# Real time selective sequencing using nanopore technology

**DOI:** 10.1101/038760

**Authors:** Matthew Loose, Sunir Malla, Michael Stout

## Abstract

The Oxford Nanopore MinION is a portable real time sequencing device which functions by sensing the change in current flow through a nanopore as DNA passes through it. These current values can be streamed in real time from individual nanopores as DNA molecules traverse them. Furthermore, the technology enables individual DNA molecules to be rejected on demand by reversing the voltage across specific channels. In theory, combining these features enables selection of individual DNA molecules for sequencing from a pool, an approach called ‘Read Until’. Here we apply dynamic time warping to match short query current traces to references, demonstrating selection of specific regions of small genomes, individual amplicons from a group of targets, or normalisation of amplicons in a set. This is the first demonstration of direct selection of specific DNA molecules in real time whilst sequencing on any device and enables many novel uses for the MinION.

---

The introduction of nanopore sequencing, represented commercially by Oxford Nanopore Technologies (ONT) MinION sequencer, has enabled a new sequencing paradigm for real time analysis^1^. MinION reads are collections of electrical current measurements, known as squiggles, where the values are determined by the specific DNA bases in contact with the nanopore at a given instance in time (Fig 1A). Unlike all other sequencing platforms, the sequencer is able to stream current values from individual nanopores as DNA molecules are passing through, enabling the analysis of current traces whilst sequencing is in progress. The MinION can stream data from all channels (currently 512) simultaneously, with each channel individually addressable and able to reverse the voltage across its pore, thus rejecting the read. In principle this enables selective sequencing. Molecules can be sampled by the sequencer and, based on some criterion, either be allowed to sequence to completion or be rejected and a second molecule sampled. On rejection the read would be ejected from the nanopore back into the pool of molecules. This molecule is not likely to be available for sequencing again as the enzymes associated with the leader sequence will already have migrated along the sequence. Furthermore the strength of rejection is sufficient that the DNA is physically remote from the channel. This type of approach has been described, but not yet implemented, by Oxford Nanopore and has been named ‘Read Until’.

**Figure 1.**
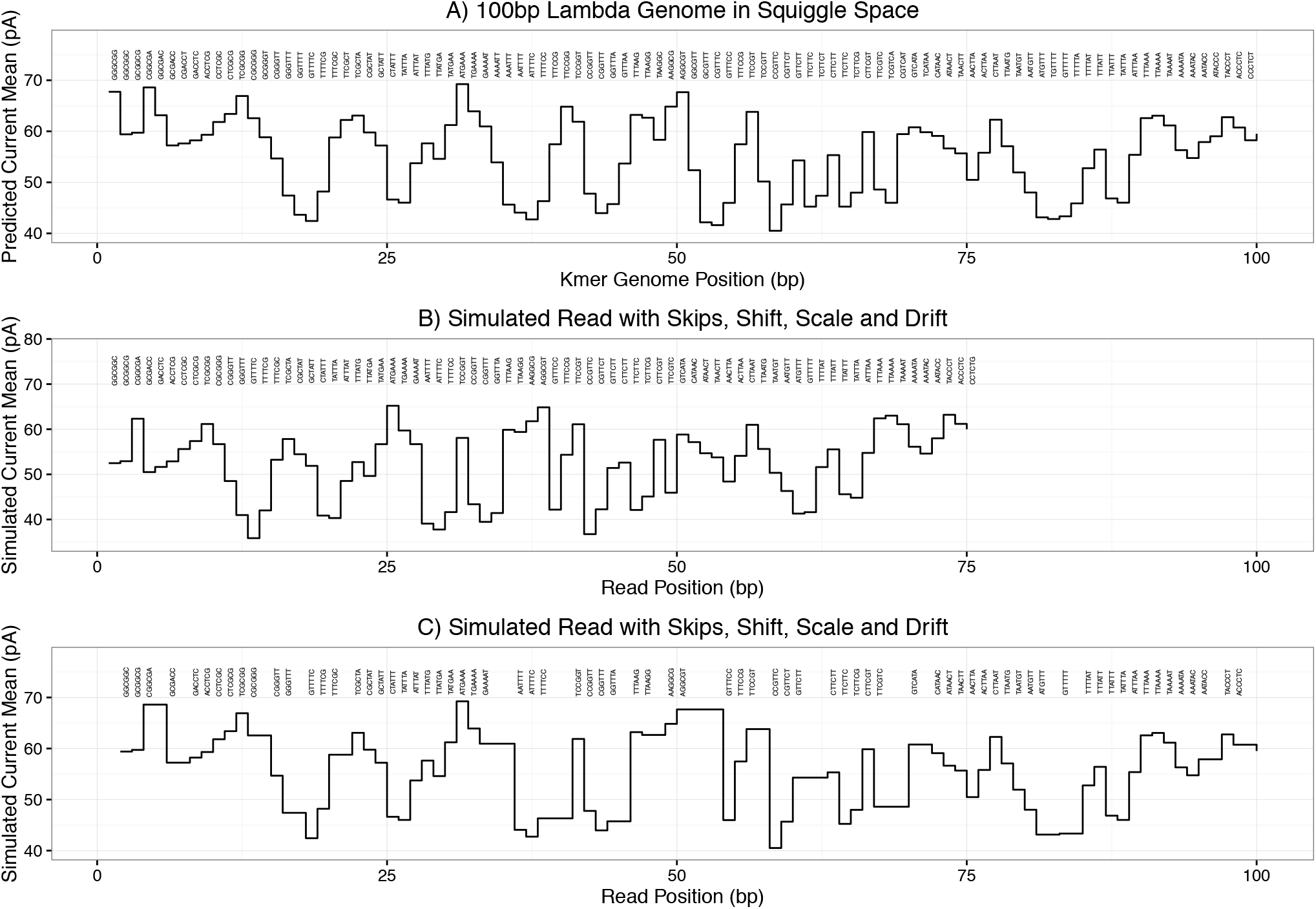
- Simulated reference and reads in squiggle space. Panel A shows a model squiggle inferred from the first 100 bp of bacteriophage lambda. Individual kmers are shown above each event in the squiggle. Panel B shows an example read derived from the same 100 bp region as in A but incorporating shift, scale and drift, along with randomly skipped kmers. Panel C shows this same read, but stretched in the time axis to map directly to the original reference.

To maximize the benefit of selective sequencing identification must be rapid and accomplished before the read is complete. This depends on two key parameters. Firstly, the speed of sequence identification and placement with respect to the reference and secondly, the average length of reads within the library. Read length is limited by the ability to generate the input material, with mappable reads reported exceeding 100 kb^2^. Fully exploiting ‘Read Until’ requires matching the shortest possible fragment of the longest possible read to a specific position within a reference sequence. However, implementation of ‘Read Until’ is complicated by the fact that the sequence data streamed by the device is a current squiggle, not a decoded sequence of known bases. To date there are no publicly available methods for analysing squiggle data directly without first base calling.

One possible approach would be to base call these read fragments, but all current implementations of base calling are too slow for this purpose and benefit from subsequent optimisations to maximise base call quality^1,3-5^. Therefore, we focused on mapping squiggle data directly to a reference sequence in squiggle space. The mapping of one squiggle to another shares much in common with the comparison of audio signals and so we turned to dynamic time warping (DTW), an algorithm first applied to the matching of speech^6,7^. Guaranteed to find the optimal alignment of two series of time-ordered data, DTW has been used many times in the analysis of sequence data, recent examples including alignment of genomic signals^8^. Here we use DTW to match observed short squiggles to a reference in real time, manipulating the output from the sequencer to demonstrate selective sequencing on the MinION. We apply this approach to two problems. First, the selection of specific regions from a whole genome library, in this case Lambda. Second, the selective sequencing of individual amplicons from a pooled set of molecules to provide efficient coverage balancing, motivated by recent field experiments with Ebola^9^. All methods and scripts are freely available under the MIT license from https://github.com/mattloose/RUscripts and we also include methods for post processing reads after sequencing is complete. Figures are presented as Jupyter notebooks at https://github.com/mattloose/RUFigs. All read data are available from the ENA (PRJEB12567). The API required for addressing the sequencer is available directly from Oxford Nanopore Technologies on request but is currently an alpha release and still in development.

## Squiggle Matching nanopore reads

To match short fragments of read data to a reference requires the conversion of the reference sequence to a hypothetical current trace. In brief, the appropriate model can be extracted from a base called MinION read, and thus an expected squiggle can be calculated for a reference sequence (Fig 1A) (see methods). Given two perfect squiggles it is theoretically possible to match one to another by simple visual inspection. This process is complicated by the changing dynamics of the sequencing environment including voltage changes, noise and interactions between channels. These changing behaviours can be captured by three variables, shift, scale and drift (Fig S1A). The base caller provided by ONT most likely calculates these values using expectation-maximisation but derives the data to do so from the entire read^5^. For short nanopore reads z-score normalisation can be used which overcomes shift and scale, but not drift (Fig S1B). Based on examination of these parameters in base called reads, we concluded that drift is not a significant problem and so Z-score normalization can be used with both reference and incoming read fragments.

Nanopore reads also have a low frequency of errors in event detection such that reads contain event insertions and deletions as well as small time differences (dwells) when translocating identical DNA molecules (compare Fig1A-C)^5^. To compensate for the residual differences in signal between the read and modelled reference, the shift in time and potential insertions/deletions in events, we use DTW subsequence search. This algorithm is able to measure the similarity between two temporal sequences which vary in amplitude or speed and has been used in many problems involving temporal sequence recognition^7,8^. Thus we can determine the optimal mapping position of the query sequence to both template and complement reference squiggle and use the distance score to determine the optimal mapping (see methods).

We tested this approach on synthetic data to determine our ability to uniquely match a sequence of arbitrary length within a reference. A parameter scan shows any 10mer squiggle from the lambda reference can be correctly mapped with DTW (Fig S2A). On real data longer query sequences are required presumably due to the inherent noise. In the absence of normalisation, less than 40% of real data can be mapped correctly within 50 bp of the known reference position (Fig S2B). This can be dramatically improved by z-score normalisation for real data (Fig S2 C,D). It is well known that the naive DTW algorithm has a complexity of O(m*n) where here m is the reference length and n is the query length. Reducing the query sequence length will therefore improve the search time (Fig S2). We chose 250 events as a conservative balance between search speed and accuracy (see methods). At current sequencing speeds (70 b/s) 250 events represents approximately 3.5 seconds of data collection, although some of the data presented here are derived from the slower SQK5 data at 30 b/s.

## Target Enrichment

To test ‘Read Until’ functionality with the MinION, we developed two applications using bacteriophage lambda DNA; target enrichment and amplicon sequencing. The choice of lambda is favourable as the small genome size (48,502 bp) results in a limited total search space of 97,004 bp. For our first experiment, we enriched for two 5kb regions of the genome, from 10-15 kb and 30-35 kb, rejecting all other reads. As an internal control, we exploited the ability to address each channel of the MinION flow cell individually, only applying ‘Read Until’ on even numbered channels. Adjacent channels are sampling reads from the same pool of molecules allowing for a direct comparison. We ran the sequencer for 47 hours generating 31,609 reads of which 8,901 were 2D. This gave a mean coverage of 840X for 2D reads (i.e both template and complement strands have been sequenced for a single read) across the lambda genome (using the slower SQK5 chemistry). Looking at all reads regardless of channel, two peaks are observable covering the two regions of enrichment (Fig 2A black dashed line). Splitting the data into odd and even numbered channels reveals even numbered channels rejecting reads mapping outside the region of interest (Fig 2A red line). In contrast, reads originating from the entire genome are found within the odd numbered channels (Fig 2A blue line). The spread of the peaks observable in Fig 2A correlates well with the mean length of 2D reads in this particular library at 5kb. Repeating this experiment using the faster 70 b/s SQK6 chemistry, selecting for reads from 10-15kb and 30-35kb, shows the same result (Fig 2B). This experiment generated 48,195 reads in 18 hours. To validate our ability to be selective we then switched approach on this specific flowcell and library, running ‘Read Until’ on all channels selecting those reads mapping within a window of 15-25 kb (Fig 2C). This illustrates one use for ‘Read Until’: regions for sequencing can be prioritised and once a specific coverage depth or goal has been achieved, a second experiment can be conducted analysing an alternative region from the same library. To process the data from the flowcell fast enough these experiments were run using 22 cores (Intel(R) Xeon(R) CPU E5-2690 v3 @ 2.60GHz) on a separate server from that running the MinION device.

**Figure 2.**
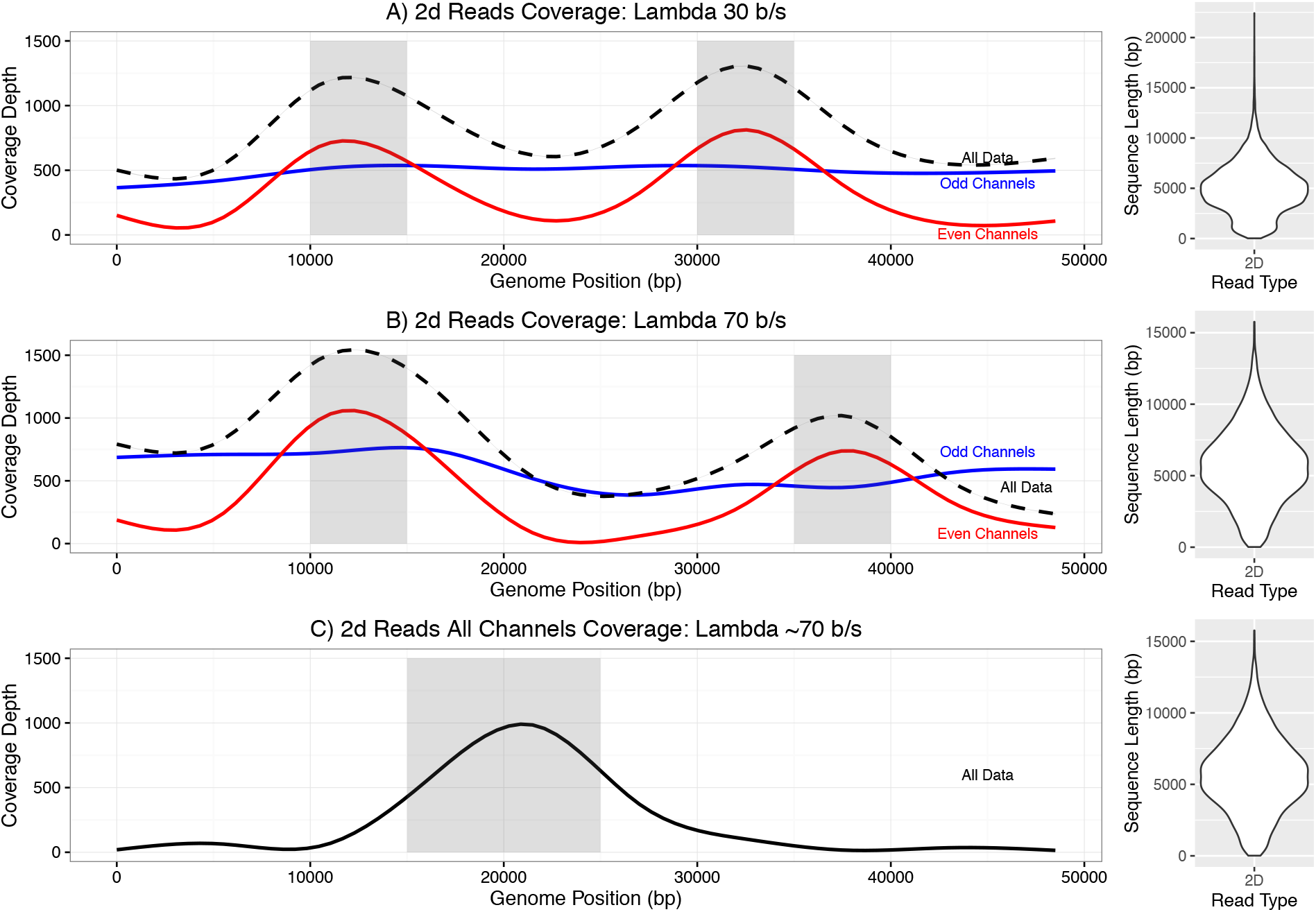
- Selective enrichment of targeted regions of the lambda genome. Panel A shows selective enrichment of the lambda genome in two 5kb regions (10-15 kb and 30-35 kb) sequencing with SQK5 chemistry (30b/s). Read Until is only applied to even numbered channels. (All Channels-black dashed line, odd numbered channels - blue line, even numbered channels - red line) Panel B shows selective sequencing using SQK6 chemistry (70 b/s) enriching at 10-15 kb and 35-40 kb. Read Until is only applied to even numbered channels. Colours as in A. Panel C shows selective sequencing on all channels of one 10 kb region (15-25 kb). Violin plots show 2D read length for each library.

## Amplicon Sequencing

The speed of match obtainable by DTW is inherently constrained by the length of the reference as the algorithm is of O(m*n) complexity^6^. We thus changed problems to consider amplicon sequencing on the MinION device, for which a smaller problem space can be defined. This is an approach to which the MinION in its current form is ideally suited. Indeed, the first major field study to use the MinION exploited such an approach to rapidly sequence amplicons derived from the Ebola genome^9^. For phylogenetic analysis of the Ebola genome, Quick and colleagues derived an amplicon based strategy to sequence the approximately 19kb Ebola genome using either 11 or 19 amplicons in the field. A challenge for any such field application is to ensure adequate coverage of every amplicon with as little total sequencing as necessary to minimise the total resource required and the volume of data to transfer for subsequent analysis.

Several optimisations are possible for amplicon matching. Firstly, sequence that is not from the start or end of an amplicon can be removed from the reference, resulting in a shorter reference squiggle and a concomitant increase in the speed of matching. Given that an amplicon can only be sequenced from a 5’ end, the effective reference space can be halved by removing unnecessary complement sequence. For example, a 900 bp window around the anticipated start site for an 11 amplicon approach reduces the reference to 19,800 rather than the 38,000 events required for mapping to the whole Ebola genome. Scaling this approach up to larger genomes becomes even more beneficial as the search space is dictated by amplicon number, not genome size. This reduction in search space enables the ‘Read Until’ algorithm to run on the same computer as the MinION device itself, an important point for sequencing in remote locations, without large computational resources.

Practically sequencing Ebola in a lab environment is limited by the availability of safely analysable material. We therefore evaluated the feasibility of this strategy using 11 amplicons spanning 22kb of the bacteriophage lambda genome (see methods). Individual amplicons were pooled prior to library preparation at approximately equimolar concentrations and a sequencing library prepared following the standard protocol used by Quick et al^1,9^. We then performed multiple short duration runs on a single flow cell to demonstrate the ability to select for specific amplicons. Fig 3A shows a 20 minute run without ‘Read Until’ applied sequencing from 209 channels, generating 3,657 reads of which 1,671 were 2D. All 11 amplicons are present within the sequenced sample although at different levels. We then stopped and restarted this run, applying ‘Read Until’ on all channels to exclude the even numbered amplicons from the sequencer (Fig 3B). This was successful although some leakage was observed on the rejected amplicons as had been seen with the whole genome selection. We then started a third run using the same library and flowcell selecting the inverse amplicons and successfully blocked the sequencing of odd numbered amplicons (Fig 3C). To understand the misplaced reads in more depth, we reprocessed them using the same DTW analysis to sort the reads into individual amplicon groups. Alignment of the base called data to the reference shows that the sorting is highly accurate, although a small proportion of reads are misplaced consistent with our earlier analysis of the ability to map reads (Fig S3A,B, Fig S2).

**Figure 3.**
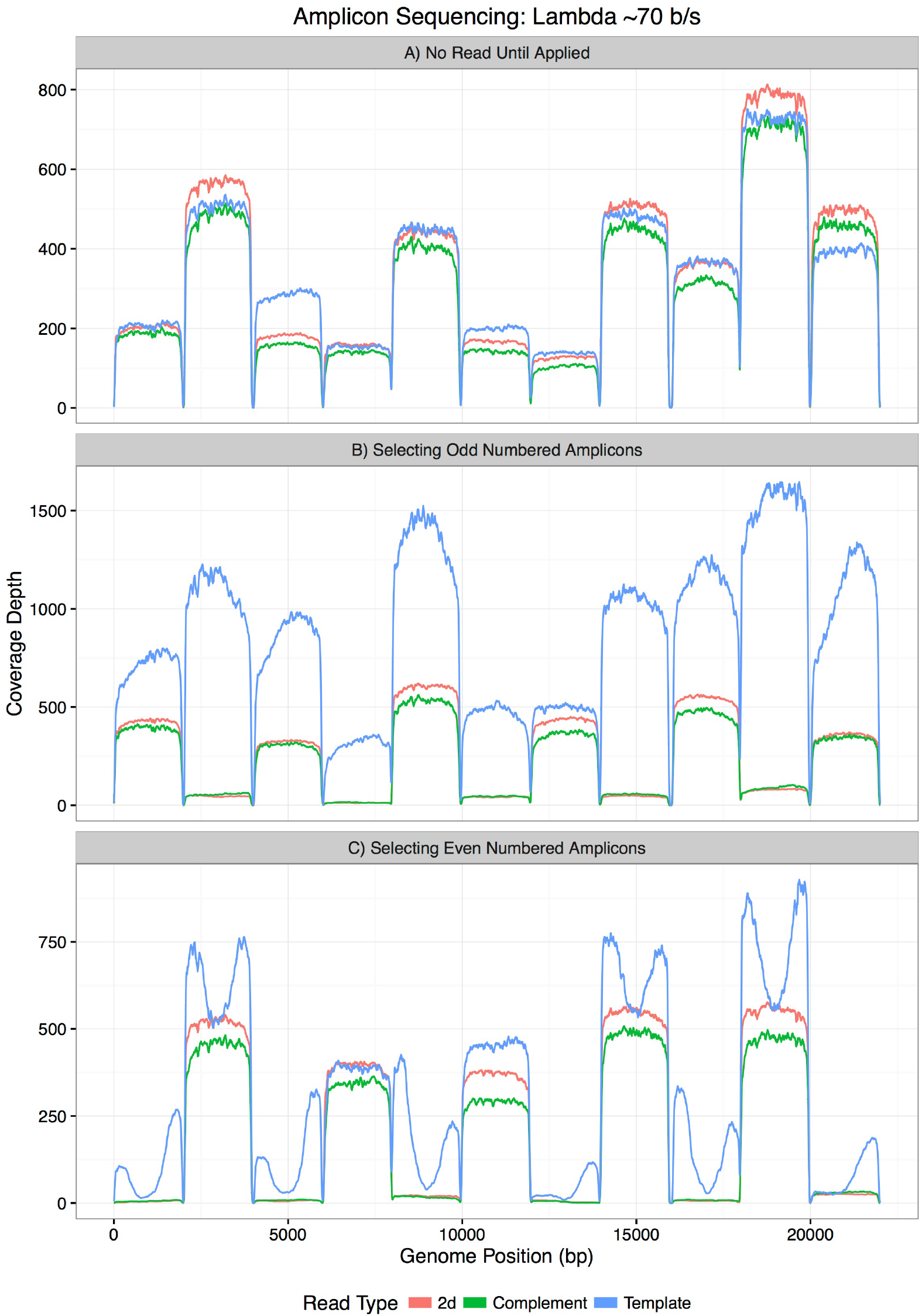
- Selective sequencing of amplicons. Panel A (Top) shows all amplicons sequenced without Read Until applied. 2D reads are pale red, complement reads are green, template reads are blue. Panel B (Middle) shows selective sequencing of odd numbered amplicons. 2D reads are absent from even numbered amplicons. Template reads can be seen for all amplicons. Panel C (Bottom) shows the inverse relationship where even numbered amplicons are selected for sequencing.

An obvious application for this technology is ensuring uniform or minimal coverage over each amplicon. The challenge here is read counting and overcoming numerous potential sources of latency within the system. From observation, the MinION device generates a number of read starts which do not result in read data being written to disk. These may represent short reads, or perhaps non DNA material blocking a pore. In addition, a read which has been mapped at the start of the template sequence may not include a hairpin sequence and so result in only a template and not a 2D read. Thus it is not sufficient to keep track of the observed reads direct from the sequencer, it is also necessary to monitor the reads as they are written to disk. This could be done by analysing base called reads but this is subject to delays in read processing by Metrichor and access to the base calling service at the time of sequencing. We therefore keep track of reads from the initial signal from the sequencer through to being written to the hard disk. An additional problem became apparent as the current version of minKnow (0.50.2.15), the software controlling the MinION, writes reads to disk after some period of buffering. This lag introduces a further delay between a read completing its translocation through the pore and the data being available on disk for analysis.

To determine our ability to normalise an amplicon library with ‘Read Until’ in real time we attempted to obtain uniform 2D coverage of our 11 amplicon library. Fig 4A shows the unnormalised library sequenced for approximately 10 minutes with differential coverage of each amplicon. We then restarted the run, introducing normalisation and rejecting reads once they had crossed a specified threshold. Our goal was to ensure at least 200x coverage of each amplicon. The Read Until script would stop the sequencer once it had detected these coverage levels. To reach this goal, the sequencer ran for 48 minutes and the relative normalisation can be seen in Fig 4A. Once base called, the 2D coverage is much greater than 200x. To see if this was consistent with latency within the system rather than leakiness in blocking of individual amplicons we investigated mean coverage depth in 1 minute intervals over the run (Fig 4B, Fig S4). This clearly reveals the time at which the sequencer begins to reject each amplicon. Those with the highest coverage in the library are rejected first (amplicons 2,3,5,8,10 and 11), whilst those with lowest coverage continue sequencing until the end of the run (see amplicons 4 and 7). Fig 4B shows examples of the dynamics observed for each amplicon type. Amplicon 3 accumulates 2D reads for the first 20 minutes of the run, but from that point reads are rejected. Amplicon 4 accumulates 2D reads at a steady rate until 20 minutes in to the run at which point the rate of its sequencing increases. This is due to higher sequencing capacity being directed towards Amplicon 4 as other amplicons sequencing coverage is reached (∽350x 2D sequencing). Amplicon 6 sees a similar trajectory although it reaches its peak after around 35 minutes and then all subsequent reads are rejected.

**Figure 4.**
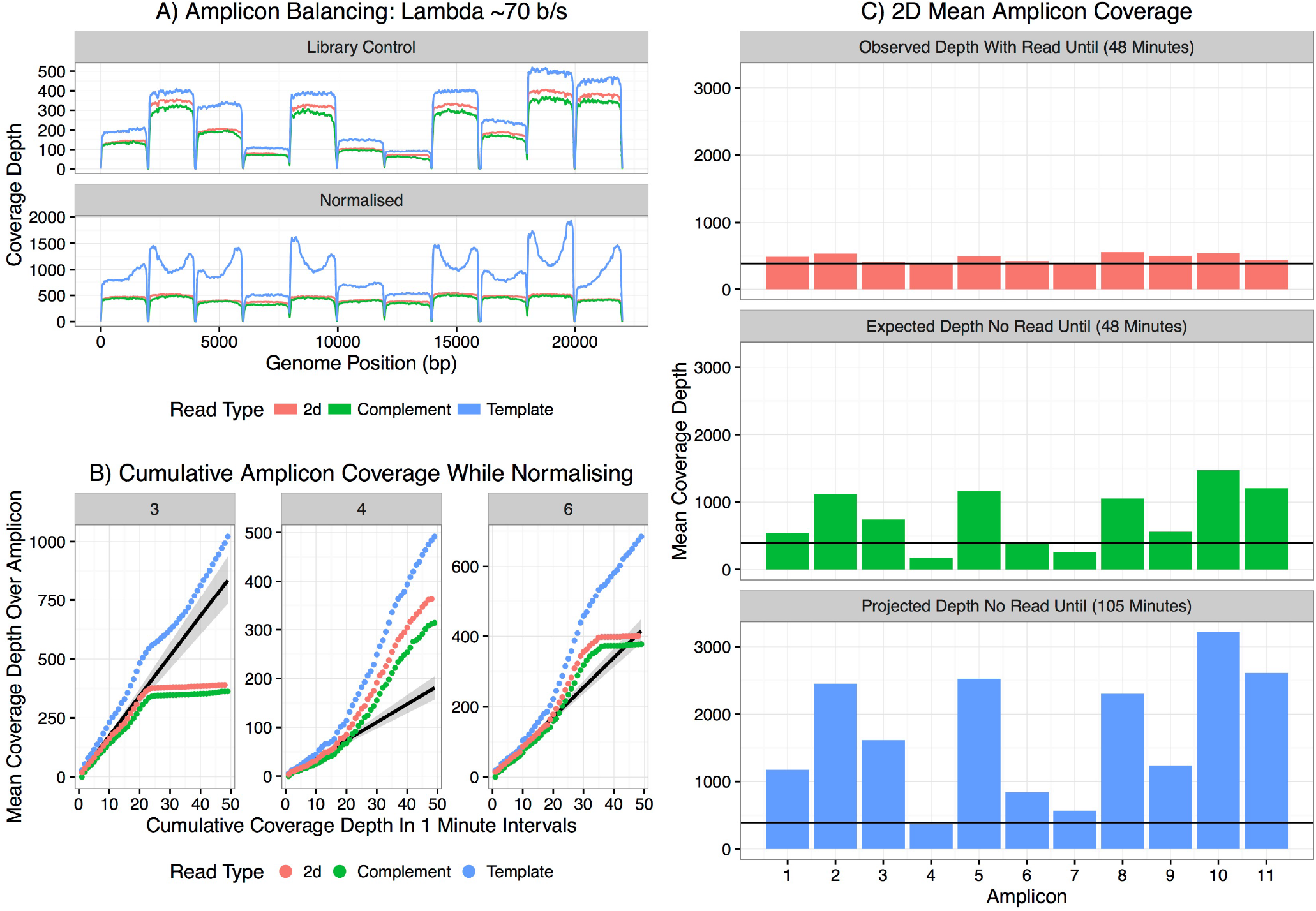
- Normalisation of amplicons using ‘Read Until’. Panel A shows coverage over each of the 11 amplicons without ‘Read Until’ (top pane) and with ‘Read Until’ applied (bottom pane) trying to normalise coverage. Pale red - 2D reads, Green - complement, Blue - Template. Panel B shows the cumulative coverage for three representative amplicons in 1 minute intervals. The black line is predicted 2D coverage for each amplicon calculated by fitting to the first 15 minutes of data. Panel C shows the observed 2D coverage after 48 minutes (top pane), the total coverage predicted for each amplicon based on the first 15 minutes of sequencing (middle pane) and the projected depth if the reads were allowed to accumulate without ‘Read Until’ to match the minimum coverage seen with ‘Read Until’ which would take 105 minutes (bottom pane).

By extrapolation from the first 15 minutes of data collected during this run it is possible to compare the observed levels of coverage, with the expected coverage depth after 48 minutes without ‘Read Until’ applied (Fig 4C). The horizontal line in these plots indicates the threshold for coverage depth at which the run completed (including the latency previously described). Similarly we can show the mean depth of coverage we would expect to see in order to have all amplicons at this minimal level. This would have taken 105 minutes to achieve and generated thousands of unnecessary reads slowing the rate of subsequent analysis.

To our knowledge this is the first demonstration of selective sequencing of specific molecules without prior enrichment during sample preparation. This is also the first demonstration of the ability to map sequence reads from a nanopore device without first converting the signal into base sequence. The specific method we describe here, a basic application of DTW from the mlpy library, is surprisingly accurate. Although the current implementation is not sufficiently fast for large genomes, the algorithm has the potential to be optimized both algorithmically, using a number of well described approaches including lower bounding, and computationally by implementation on GPU or FPGA processors^10,11^.

Notwithstanding the challenges of searching through large genome space, we demonstrate that this technique can be applied to amplicon sequencing. Here the experimental design can be used to optimise the search space such that reads only need to be mapped to a subset of the reference genome. The ability of nanopore technology to sequence longer amplicons could be further exploited. A clear advantage of this approach is in viral outbreaks where rapid sequencing can help in tracking and characterisation of the emerging outbreak. The extreme portability of the MinION sequencer allows it to be easily deployed to remote locations. However, these locations may have restricted internet access and so cloud based base calling may be a challenge. ‘Read Until’ methods can also be applied post sequencing to sort and pre-select reads to be base called using limited bandwidth connections.

Exploitation of the real time nature of the minION platform, where reads are available for analysis as soon as they have passed through the sequencer, has been shown by many groups^12-14^. Here we demonstrate true real time analysis and the potential for selective sequencing on a MinION device with ‘Read Until’. The anticipated increasing speed of nanopore sequencing (’fast mode’) and the scaling up of the MinION to 3,000 channels, and the PromethION with 144,000 channels, will challenge the implementation of ‘Read Until’ in real time and require algorithmic enhancements and computational power. Yet ‘Read Until’ based approaches will enable new approaches to sequencing in the future, such as exonic sequencing without pre-sequencing target capture, controlled depth of coverage over entire genomes, novel counting applications for RNA seq and many applications that have yet to be imagined.

## Author Contributions

ML conceived and designed the project, implemented the code, ran the sequencing, performed analysis and wrote the manuscript, MS carried out analysis, SM generated libraries and ran the sequencing.

## Acknowledgements

The authors thank Nick Loman and Josh Quick for helpful discussions, advice and motivation, Teri Evans for comments on the manuscript and figures and Martin Blythe for discussions. We also thank Oxford Nanopore Technologies for access to the Read Until API and helpful discussions and support.

